# Inhibition of miR-214 expression by small molecules alleviates head and neck cancer metastasis by targeting ALCAM/TFAP2 signaling

**DOI:** 10.1101/2023.04.04.535560

**Authors:** Anshu Agarwal, Vikash Kansal, Humaira Farooqi, Ram Prasad, Vijay Kumar Singh

**Affiliations:** Department of Zoology, Agra College, Dr. B. R. Ambedkar University, Agra-282004 (India); Department of Otolaryngology, Emory University, Atlanta, GA 30322 (USA); Department of Biochemistry, Hamdard University, New Delhi-110062 (India); Department of Ophthalmology and Visual Sciences, University of Alabama at Birmingham, Birmingham, AL-35294 (USA); Narain PG Degree College, Shikohabad, Dr. B. R. Ambedkar University, Agra-282004 (India)

**Keywords:** miR-214, invasion, metastasis, ALCAM, TFAP2, head and neck cancer

## Abstract

Predominantly, head and neck cancer (HNC) is considered a regional disease and develops in the nasal cavity, oral cavity, tongue, pharynx, and larynx. In the advanced stage, the HNC spread into distant organs. By the time head and neck cancer diagnosed, the estimated metastasis is occurred in 10-40% cases. The most important vital organs affected by distant metastasis are the lungs, bones, and liver. Despite several advancements in chemotherapies, no significant changes are observed as 5-year survival rate remains the same. Therefore, it is crucial to decipher molecular mechanisms contributing to the metastatic dissemination of head and neck cancer. Here, we tested a novel ALCAM/TFAP2 signaling by targeting multidisciplinary miR-214 expression in head and cancer cells. Our results revealed that HNC cell lines (CAL27, SCC-9, SCC-4, and SCC-25) exhibit higher expression of miR-214 compared with normal human bronchial epithelial (NHBE) cells. Higher expression of miR-214 drives the invasive potential of these cell lines. Down-regulation of miR-214 in CAL27 and SCC-9 cells either using an anti-miR-214 inhibitor (50nM) or a small molecule of green tea (EGCG) inhibited cell invasion. Treating CAL27 and SCC-9 cells with EGCG also reduces ALCAM expression, a key activated leukocyte cell adhesion molecule, potentially blocking mesenchymal phenotype. Dietary administration of EGCG significantly inhibits distant metastasis of SCC-9 cells into the lungs, liver, and kidneys. Our results also demonstrate that the reduction of miR-214 expression influences in vitro cell movement and extravasation, as evident by reduced CD31 expression, a neovascularization marker. Together, these studies suggest that identifying bioactive molecules that can inhibit distant metastasis regulated by the miRNAs may provide potent interventional approaches and a better understanding of the complex functions of miRNAs and their therapeutic targets for clinical application.

## 1. Introduction

Despite its origin, treating any cancer, including head and neck cancer, is always challenging. Current therapeutic regimens do not provide effective measures, as there is no significant improvement in the five-year survival rate. The term head and neck squamous cell carcinoma (HNSCC) represents all anatomical sites of the head and neck region, emphasizing the larynx, pharynx, and oral cavity. The annual global incidence of HNSCC has been estimated to be approximately 660,000 new cases and about 325,000 deaths in 2023[1]. In general, men are at greater risk than women[2]. Although HNSCC is predominantly considered regional cancer, its spread into the other organs is also a major determinant of its management and diagnosis. Although in comparison to other malignancies, the incidence of HNSCC metastasis is low ∼15%[3,4]. However, if distant metastases of HNSCC occur, the patient does not survive long, is considered incurable, and is treated in a palliative manner[5]. The lungs, bone, liver, brain, and skin are the most common sites of HNSCC metastasis[3,6]. The distant metastasis of HNSCC is a complex mechanism governed by various genetic, epigenetic, and phenotypic changes. The conversion from the regional to metastatic stage of HNSCC is a series of well-characterized and defined molecular changes, which involve alterations in gene expression associated with adhesion, invasion, migration, extravasation, epithelial to mesenchymal transition, and angiogenesis.

The role of microRNAs (miRNAs) in regulating HNSCC growth, cell differentiation, and migration/invasion is now being recognized as key regulators. The miRs are small, approximately 18-25 nucleotides long, endogenous, and highly conserved non-coding RNA molecules. These miRNAs regulate gene expression by binding with 3’-untranslated regions (3’UTRs) of target mRNAs, which have complete or partial complementarity with their seed region. This leads to either mRNA destabilization or the inhibition of translation initiation, or both[7-10]. Studies demonstrated that miRs are largely involved in various cellular processes such as cell proliferation, growth, differentiation, metastasis, and extravasation through the fine-tuning and alterations in signaling pathways and contribute to cancer progression[11]. The differentially dysregulated expression of miRs is correlated with cancer development, as many of these miRs possess oncogenic or tumor suppressor activity in various tumors[12,13]. The increased expression of miR-214 has been observed in several cancers, such as melanoma, ovarian, and gastric cancer, while low expression in breast cancer[14,15]. Studies have shown that the involvement of miR-214 is not only limited to cancers of different organs but also plays important roles in pulmonary arterial hypertension and myocardial apoptosis[16,17]. Recently, It is has been reported that over-expression of miR-214 in melanoma facilitates its progression through activated leukocyte cell adhesion molecule (ALCAM) and transcription factor activated protein-2 (TFAP-2)[18]. The over-expression of ALCAM is reported in many tumors and is considered a prognostic molecular marker[19], while loss of TFAP-2 expression is reported in other neoplasms [20]. TFAP-2 is a family of transcription factors characterized to regulate the developmental process between growth and differentiation[21,22].

The naturally bioactive phytochemicals such as Epigallocatechin gallate (EGCG), an active component of green tea, are emerging as new options for better management and as preventive and interventional regimens for a different types of cancers with greater efficacy and least or minimum toxicity. To better understand the molecular mechanisms underlying distant metastasis of HNSCC, our study aimed to reveal the prognostic and therapeutic values of miR-214 to suppress metastatic HNSCC. Here, we have shown the anti-metastatic effect of EGCG in terms of the invasive potential of HNSCC cells and whether it is mediated through their effect on miR-214 expression. For this purpose, we used CAL27, SCC-9, SCC-4, and SCC-25 cell lines as an *in-vitro* model and ascertained whether EGCG inhibits their metastatic potential through its inhibitory effect on miR-214 expression. Further, we present evidence that EGCG inhibits metastasis of SCC-9 cells *in-vivo* mouse metastatic model and that they do so through: (i) down-regulation of miR-214 expression and (ii) blocking of invasion/extravasation of HNSCC through inhibition of ALCAM and restoration of TFAP-2 expression in lung, liver, kidney and spleen tissues.

## 2. Materials and Methods

### 2.1. Cell lines, culture conditions, and treatment

The HNSCC cell lines: CAL27 (#CRL-2095), SCC-9 (#CRL-1629), SCC-4 (#CRL-1624), and SCC-25 (#CRL-1628), and normal bronchial epithelial cells (NHBE) (#PCS-300-010) were purchased from ATCC (Manassas, VA). The cancer cells were cultured as monolayers in Dulbecco’s modified Eagle’s medium (DMEM). The 10% heat-inactivated fetal bovine serum (FBS; v/v) was added to the cell culture medium to supply the naturally required factors for the cell’s attachment and growth. Two antibiotics, penicillin and streptomycin (100 μg/mL), were also added to the medium to protect the cells from contamination and maintain aseptic conditions. The cell cultures were maintained in an incubator with 5% CO2 at 37^0^C. The NHBE cells were cultured in NHBE specific medium.

For the *in-vitro* treatment, EGCG was dissolved in the cell culture medium to achieve the desired concentrations and treated in a dose-dependent (0, 1, 5, and 10 μg/mL) manner.

### 2.2. Chemicals, reagents, and antibodies

The purified EGCG (95%) was purchased from the commercial herb supplier, Maysar Herbals (Haryana, India). The primary antibodies specific for ALCAM (#sc-74557), CD31 (#sc-376764), β-actin(#sc-47778), Histone H3 (#sc-5661), and secondary antibodies horseradish peroxidase(HRP)-linked anti-mouse (#sc-2031), HRP-anti-rabbit (#sc-2030) from Santa Cruz Biotechnology (Santa Cruz, CA, USA), TFAP-2A (#HPA056871) and TFAP-2C(#SAB2102408) primary antibodies were purchased from Sigma-Aldrich (St. Louis, MO). Anti-miR-214 inhibitor (#4464084), lipofectamine (#11668019), and custom primers specific for miR-214 and U6 were obtained from Invitrogen (Carlsbad, CA). All antibodies and other chemicals were purchased through an Indian supplier (New Delhi, India). The cell culture mediums and fetal bovine serum (FBS) were purchased from Thermo Fisher Scientific India Pvt. Ltd., Mumbai, India.

### 2.3. Invasion assay using Boyden chamber

The invasion potential of HNSCC cell lines was determined using Boyden Chambers (Gaithersburg, MD) and matrigel-coated millipore membranes (6.5 mm diameter filters, 8 μM pore size). Briefly, cells (15×10^3^ cells/ 100 μL medium containing 0.5% FBS) were placed in the upper section of the Boyden chambers, and the test agents were added into the upper chamber (200 μL). The chambers were assembled and kept in a cell culture incubator for specified periods. After incubation, cells from the upper surface of the membranes were removed through gentle swabbing, and cells on the lower surface of the membranes were fixed in chilled methanol and stained with crystal violet. The represented photographs from 4-5 randomly selected membrane fields were taken using an Olympus microscope equipped with a color camera. Each experiment was repeated three times.

### 2.4. miR extraction and RT-PCR

Total RNAs, containing ∼95% miRs, were isolated from cultured HNSCC cell lines and NHBE using the TRIZOL-chloroform extraction procedure described previously[23,24]. Briefly, 70-80% confluent cultured cells were washed with ice-cold PBS buffer, and the cells were covered with 1 mL Trizol reagent and immediately harvested by scraping. The lysate was transferred into a 15 mL v-shaped tube, and 0.2 ml of chloroform was added for phase separation. The cell mixture was vortexed and centrifuged at 12,000 rpm for 15 min at 4^0^C. After centrifugation, the uppermost colorless layer was separated into another tube, and 0.5 mL of isopropyl alcohol was added to precipitation RNAs. The samples were incubated at room temperature for 10 min and then centrifuged at 4^0^C. After centrifugation, the RNA pellets were washed with 75% ethanol, air-dried, and resuspended into nuclease-free water. The RNA concentration was quantified by spectrophotometry and used to prepare cDNA using the iScript cDNA Synthesis Kit (Bio-RAD), followed by the manufacturer’s instructions. RT-PCR was performed using platinum Taq DNA polymerase (Invitrogen, Carlsbad, CA) with human-specific primers for miR-214: Forward primer: TCTGCCTGTCTACACTTGCTG, and miR-214 Reverse primer: TGACTGCCTGTCTGTGCCT, and U6 forward primer: CTCGCTTCGGCAGCACA, U 6 reverse primer: AACGCTTCACGAATTTGCGT, as reported by Jiang et al.,[25]. The RT-PCR conditions were as follows: Stage I: 95°C for 3 min, 53°C for 1 min, 72°C for 30 sec (2 cycles); Stage II: 95°C for 3 min, 53°C for 1 min, 72°C for 30 sec (55 cycles); Stage III: 72°C for 5 min. The PCR product was run on a 2.5% agarose gel prepared in 1x Tris-acetate EDTA buffer containing ethidium bromide and analyzed using a Gel-Doc apparatus.

### 2.5. Transient transfection of miR-214

In the functional studies, the expression of miR-214 in HNSCC cells was silenced using a pre-designed anti-miR-214 inhibitor (Thermofisher, Waltham, MA) following the manufacturer’s instructions. Briefly, 1×10^5^ cells were seeded onto 6-well culture plates. CAL27 and SCC-9 cells were transfected in a serum-free medium with the anti-miR inhibitor, or scramble control probe, at a final concentration of 50nM, using Lipofectamine 2000 (Thermofisher, Waltham, MA) following the manufacturer’s protocol. After 24 h of transfection, cells were kept in a culture medium containing 2% FBS for up to 48 h. The cells were then harvested and used in the functional and invasion assays.

### 2.6. Western blot analysis

For protein expression analysis, cellular and nuclear fractions from HNSCC cell lines and tissues were prepared using the cell fractionation kit purchased from Cell Signaling (#9038) following the manufacturer’s instructions. This cell fraction kit also contains a protease inhibitor to avoid protein degradation [26]. Proteins were resolved using 10% SDS-PAGE gels and transferred onto a nitrocellulose membrane. After blocking the non-specific binding sites, the nitrocellulose membranes were incubated with the primary antibody specific for ALCAM, TFAP2A, and TFAP2C overnight at 4^0^C, followed by incubation with secondary HRP conjugated antibodies. Specific protein bands were visualized using the enhanced chemiluminescence reagents. Equal loading of proteins on the gel was verified by stripping the membrane and re-probing with an anti-β-actin and histone H3 antibodies.

### 2.7. Target prediction of miR-214: In silico analysis

To perform the target prediction of miR-214 and its binding with genes associated with cell migration/invasion, a web-based database (https://www.targetscan.org) was approached. Based on the analyzed sequence, this database provides background information about miRs and their seed region for binding with their target genes.

### 2.8. *Generation of* luciferase reporter HNSCC cells

To facilitate the detection of experimental metastasis *in-vivo*, SCC-9 cells were transduced with a vesicular stomatitis virus G envelope (VSV-G) pseudotyped lentiviral vector for constitutive expression of both firefly luciferase and enhanced green fluorescence protein (EGFP). The vector comprised the mouse CMV promoter, followed by firefly luciferase, the encephalomyocarditis internal ribosomal entry site (IRES), a puromycin resistance gene, and the enhanced green fluorescence protein (EGFP), wherein puromycin and EGFP were fused in-frame at their 3’ and 5’ ends, respectively, with the “self-cleaving” T2A peptide-coding sequence (CMV-luciferase-IRES-puro.T2A.EGFP)[27]. The lentiviral vector, the packaging construct, and the VSV-G plasmid DNAs were co-transfected into 293T human embryonic kidney cells to create infectious, replication defective, lentiviral vector-containing particles as described previously[28]. SCC-9 cells were transduced with the vector using a multiplicity of infections. Stable, expression-positive SCC-9 cells were selected by supplementing the culture medium with 5 μg/mL of puromycin for 5 days.

### 2.9. In-vivo metastasis of SCC-9 cells in vital organs of nude mice

Female BALB/C nude mice (4-5 weeks of age) were purchased from the National Institute of Biologicals (Noida) and housed in the institutional animal resource facility. The mice colony was maintained under a 12 h dark/12 h light cycle, a temperature of 24±2°C, and relative humidity of 50±10%. The mice were given a control AIN76A diet with or without supplementation with EGCG (0.5%, w/w) and drinking water ad libitum throughout the experiment. The Institutional Animal Care and Use Committee, under the animal protocol number-201809267, approved the animal protocol used in this study.

To determine the inhibitory potential of EGCG on HNSCC metastasis, SCC-9 cells (2.5×10^6^) constitutively expressing both luciferase and green fluorescent protein (GFP) were injected intravenously into nude mice. After four weeks, the mice were sacrificed, and vital organs (liver, lungs, spleen, and Kidneys) were harvested. The GFP/luciferase reporter SCC-9 cells in all organs were detected by bioluminescence imaging after spraying D-luciferin using the Xenogen IVIS200 imaging system. The vital fluorescent organs having metastasis were evaluated and photographed. The dietary dose of 0.5% EGCG is equivalent to 5 mg/mouse/day, considering a mouse, bodyweight 25.0 gm, eats 5 grams of chow [29]. Based on the mouse dose and formula by Nair et al., the human equivalent dose (HED) of 0.5% EGCG (W/W) will be approximately 189.18 mg/day [30].

### 2.10. In vivo extravasation assay

The extravasation of GFP-tagged SCC-9 cells was determined in the tumor, liver, lungs, spleen, and kidney tissue at the end of the experiment. After the euthanization of experimental mice, all tissues were collected and washed with chilled 0.9% saline solution following fixation in 10% buffered formalin. Paraffin-embedded sections (5 μm thick) were deparaffinized, rehydrated, and then an antigen retrieval procedure was carried out, as detailed previously[11]. After blocking the non-specific binding sites, the sections were incubated overnight with a primary antibody specific for CD31 at 4^0^C. After overnight incubation, sections were washed in PBS and incubated with goat anti-rabbit IgG labeled with red-fluorescent Alexa Fluor 594. Tissue sections were finally mounted in a vectashield mounting medium with DAPI (Vector Laboratories, Burlingame, CA). The sections were then imaged using an Olympus microscope with a color camera from 5-6 microscopic fields at 20X.

### 2.11 Statistical analysis

Data were calculated using GraphPad Prism software (San Diego, CA, USA), version 8.1. Ordinary one-way ANOVA followed by Dunnett’s multiple comparisions test and Student’s T-test assessed statistical significance between the treated and untreated groups. α=0.05.

## 3. Results

### 3.1. Expression profile of miR-214 in HNSCC cell lines

To understand the role of miR-214 in HNSCC invasion, first, we checked the expression profile of miR-214 in the metastatic HNSCC cell lines (CAL27, SCC-9, SCC-4, and SCC-25) and compared its expression level with NHBE cells using RT-PCR. As shown in **Figure 1A**, the expression level of miR-214 was higher in all four HNSCC cell lines as compared with NHBE (amplicon size 67bp). Each HNSCC cell line showed a different expression level of miR-214. We observed that the expression level of miR-214 was higher to low in the following manner: SCC-9>CAL27>SCC-25>SCC-4. The expression of miR-214 was significantly higher, almost 2 folds (p=0.01, p=0.003, p=0.009, and p=0.002, respectively) in HNSCC cell lines than NHBE as estimated by densitometry quantitation of the band intensity using ImageJ software based on band intensity ratio of miR-214 *vs*. U6 (**Figure 1B**). These results were further verified using quantitative RT-PCR (qRT-PCR); qRT-PCR data also suggests significantly higher expression of miR-214 in HNSCC cells (p=0.01, p=0.003, p=0.02, and p=0.01, respectively) compared with NHBE cells (**Figure 1C**).

**Figure 1:**
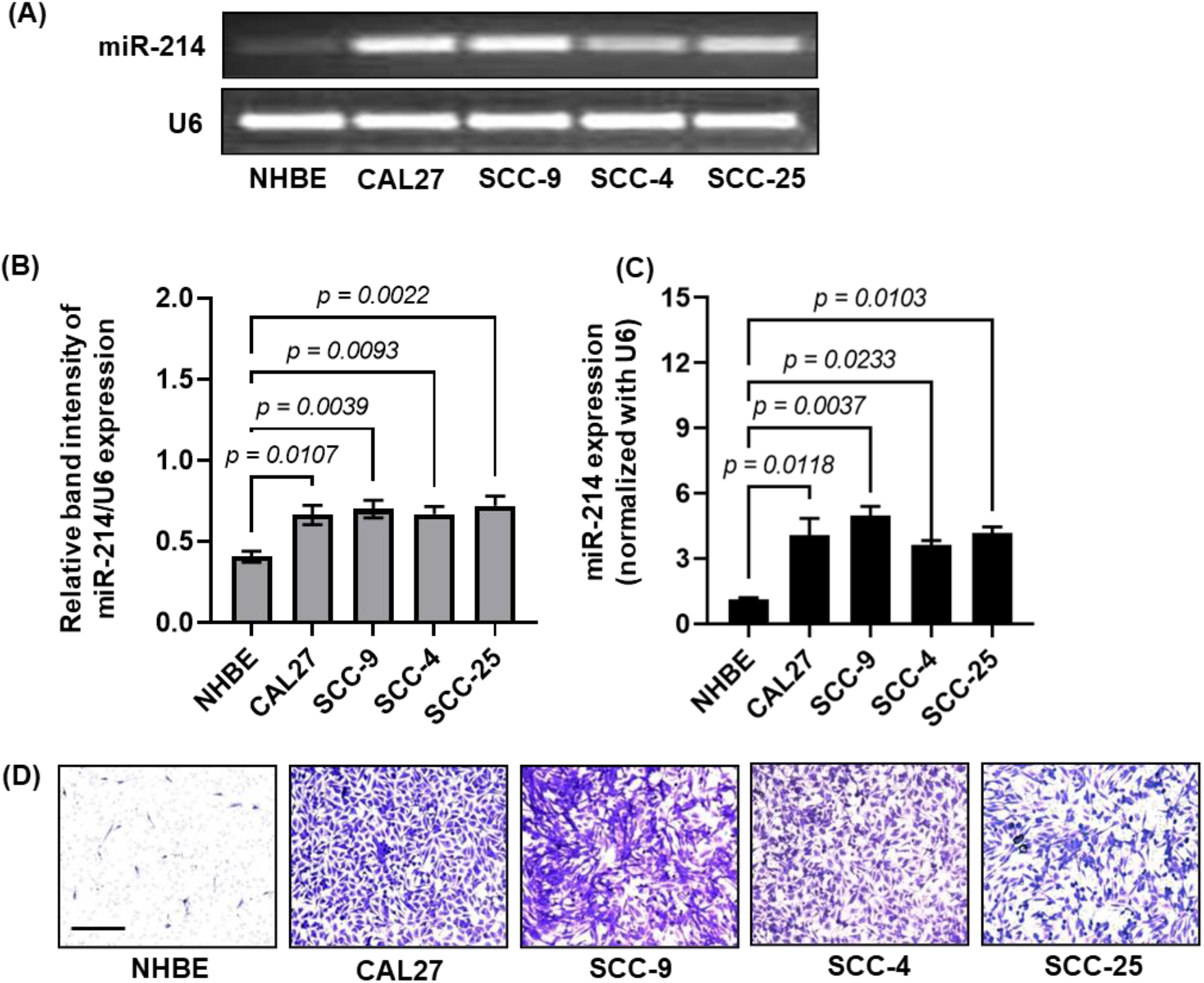
Basal levels of miR-214 expression and invasive potential of HNSCC cell lines compared with normal human bronchial epithelial (NHBE) cells. The total RNA containing miR was isolated using the Trizol-Chloroform extraction method, and cDNA was prepared following the manufacturer’s instruction using a cDNA synthesize kit (Bio-Rad). Template cDNA was used for miR-214 expression by RT-PCR **(A-B)** and qRT-PCR **(C)**. The densitometry analysis of agarose gels was carried out using ImageJ software, and data are presented as relative band intensity (gray values) mean±S.E.M, normalized with U6 **(B)**. The qRT-PCR data for miR-214 expression was normalized with U6 expression and expressed mean±S.E.M **(C)**. The invasive potential of HNSCC cell lines compared with NHBE cells **(D)**. Panels B and C were analyzed by ordinary one-way ANOVA followed by Dunnett’s multiple comparisions test. Scale bar = 20μm.

Metastatic characteristics of HNSCC make it a fatal disease with its rapid dissemination throughout the body. Next, we performed a matrigel invasion assay to determine whether miR-214 overexpression affect invasive characteristic of HNSCC cell lines. As shown in **Figure 1D**, the HNSCC cell lines overexpressed miR-214 expression, showing greater invasive potential than NHBE cells. These results indicated that miR-214 might have a role in the invasive behavior of HNSCC cell lines. Due to the greater expression of miR-214, CAL27 and SCC-9 cells were used in further studies.

### 3.2. Transient knockdown of miR-214 expression inhibits the invasive potential of HNSCC cell lines

Further, to investigate the role of miR-214 in maintaining the invasive characteristic of HNSCC cells, we knockdown the miR-214 expression in HNSCC cell lines using miR-214 inhibitor through transient transfection, and the effectivity of transient transfection was observed by qRT-PCR. As shown in **Figures 2A and 2B**, transient transfection of miR-214 inhibitor significantly decreases miR-214 expression in CAL27 and SCC-9 cell lines (p=0.01 and p=0.03). Next, we tested whether this decreased level of miR-214 affects HNSCC cell invasion. The invasive potential of HNSCC cell lines with and without knockdown of miR-214 was analyzed using a matrigel-coated membrane assay. As shown in **Figures 2C and 2D**, inhibition of miR-214 expression decreases the invasion of both cell lines. These findings showed a relationship between miR-214 and invasive characteristics of HNSCC cell lines and suggested that miR-214 can drive the invasive potential of these cell lines.

**Figure 2:**
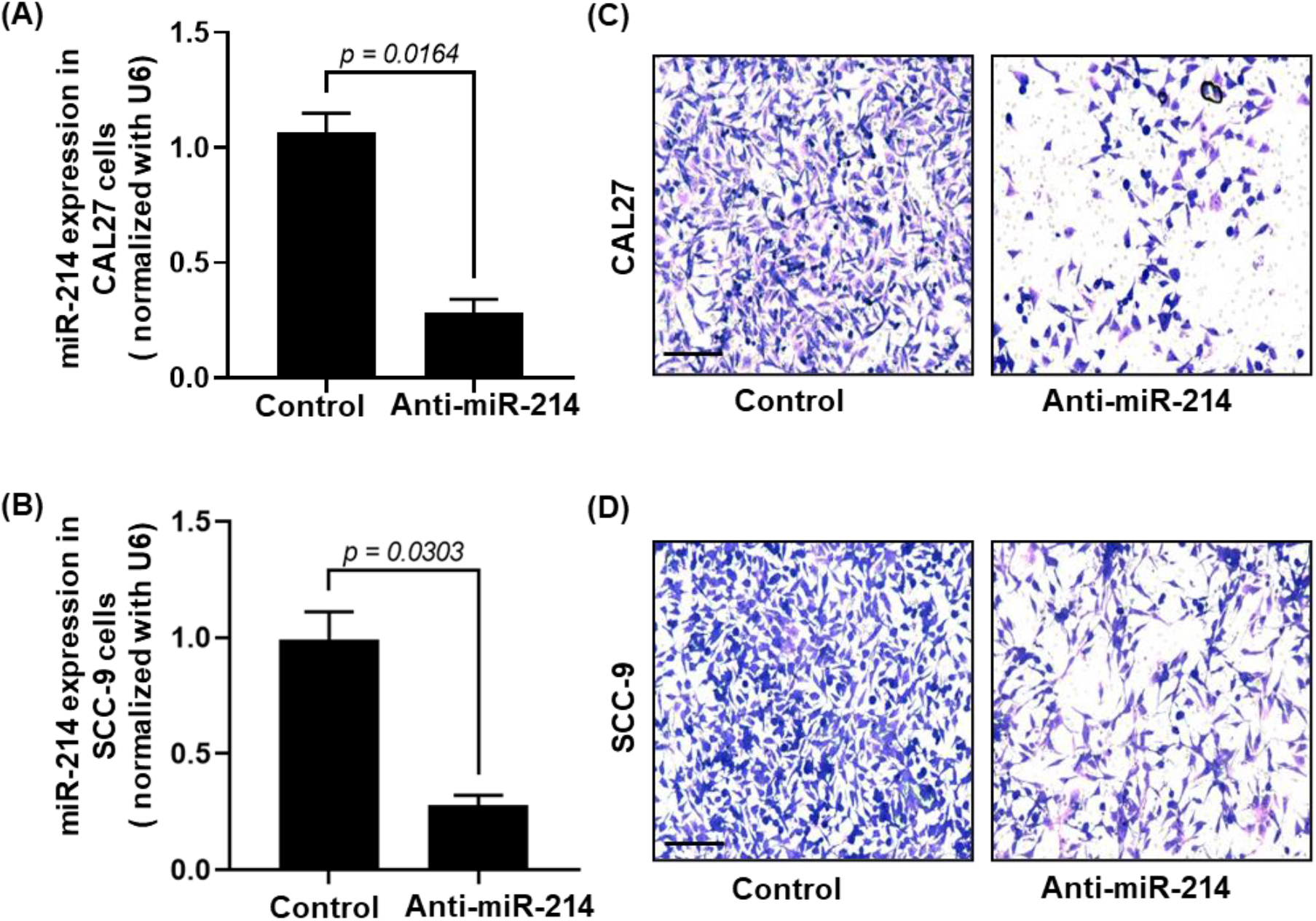
Knockdown of miR-214 inhibits the invasive potential of HNSCC cell lines. HNSCC cell lines (CAL27 and SCC-9) were transfected with anti-miR-214 (50 nM) in lipofectamine for 24 hours. After 24 hours of transfection, total RNA containing miR was isolated using the Trizol-Chloroform extraction method, and cDNA was prepared following the manufacturer’s instructions using a cDNA synthesize kit (Bio-Rad). Template cDNA was used for miR-214 expression using real-time PCR **(A-B**). Real-time PCR data were presented as mean ± S.E.M of miR-214 expression normalized with U6 compared to the non-transfected group. Effect of knockdown of miR-214 expression on invasion potential of HNSCC cell lines **(C-D)**. After transfection, the invasive potential of CAL27 **(C)** and SCC-9 **(D)** cells was analyzed using a matrigel-coated membrane in the Boyden chamber. Invaded cells were stained with crystal violet and imaged using an Olympus microscope. Panels A and B were analyzed by unpaired Student’s t-test. Scale bar = 20μm.

### 3.3. Effect of EGCG treatment on miR-214 expression and metastatic potential of CAL27 and SCC-9 cells

Bioactive molecules provide desirable health benefits besides their nutrition profile and help reduce the risk of disease development and progression. Next, to determine whether EGCG, an active component of green tea, affects miR-214 expression and metastatic potential of CAL27 and SCC-9 cells, both cell lines were treated in a dose-dependent manner (1, 5, and 10 μg/mL). To avoid the effect of cell division on migratory potential, cells were treated with EGCG for 24 hours. The estimated doubling time of CAL27 is approximately 35 hours, while SCC-9 is slowly growing and doubling in about 5-6 days. After 24 hours of EGCG treatment, the miR-214 was measured by RT-PCR. As shown in **Figures 3A and 3B**, the expression level of miR-214 was decreased in both HNSCC cell lines. This effect of EGCG on miR-214 expression in HNSCC cell lines was further verified following qRT-PCR analysis (**Figure 3C**). The expression of miR-214 was significantly lowered in CAL27 (p=0.02 and p=01, respectively) and SCC-9 (p=0.01, p=008, and p=0.004, respectively), as estimated by the relative expression of miR-214 *vs*. U6.

**Figure 3:**
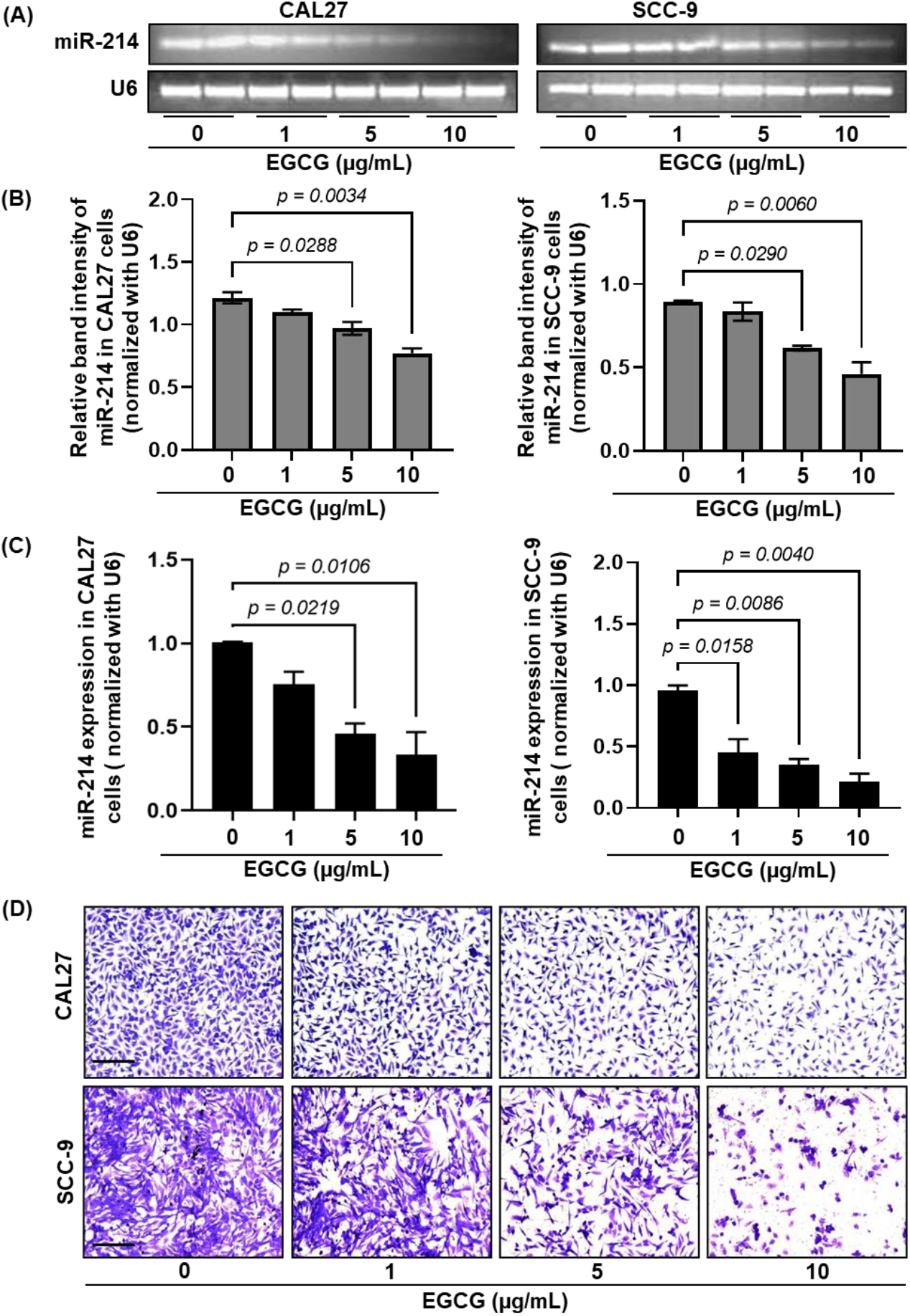
Effect of EGCG treatment on miR-14 expression and invasive potential of HNSCC cell lines. CAL27 and SCC-9 cells were treated with EGCG in a dose-dependent manner for 24 hours, and total RNA containing miRs was isolated using the Trizol-Chloroform extraction method. The cDNA was prepared following the manufacturer’s instructions using a cDNA synthesis kit (Bio-Rad). Template cDNA was used for miR-214 expression by RT-PCR **(A-B)** and qRT-PCR **(C)**. The densitometry analysis of agarose gels was carried out using ImageJ software, and data are presented as relative band intensity (gray values) mean ± S.E.M, normalized with U6 **(B)**. The qRT-PCR data were presented as mean ± S.E.M of miR-214 expression normalized with U6 compared to the untreated group **(C)**. The effect of EGCG treatment on HNSCC cell invasion potential was analyzed using a matrigel-coated transfer membrane in Boyden chambers **(D)**. Invaded cells were stained with crystal violet and imaged using an Olympus microscope. Panels B and C were analyzed by ordinary one-way ANOVA followed by Dunnett’s multiple comparisions test. Scale bar = 20μm. Each column represents an individual sample in agarose gel.

Next, the effect of EGCG was also tested on the invasive potential of CAL27 and SCC-9 cell lines. As shown in **Figure 3D**, after 24 hours of treatment, EGCG blocks the invasion of CAL27 and SCC-9 cells, as observed by the reduced number of cells on the matrigel-coated membrane.

### 3.4. The inhibitory effect of EGCG on the invasive potential of CAL27 and SCC-9 cell lines is associated with the reduction of ALCAM Expression

Cell-adhesion molecules have been reported for their significant contribution to the malignant phenotypes of human cancer cells and play an important role in cell proliferation, migration, invasion, and metastasis[31-33]. The ALCAM is a marker of mesenchymal cells and is involved in cancer metastasis[34]. Although a direct relationship between ALCAM and up-regulation of miR-214 in melanoma has been reported previously[18], HNSCC remains unexplored for this association. Therefore, to investigate whether the inhibitory effect of EGCG on HNSCC invasion is associated with decreased miR-214 mediated inhibition of ALCAM expression, we determined the levels of ALCAM in HNSCC cell lines (CAL27 and SCC-9) from the various treatment groups using western blot analysis. As shown in **Figure 4A**, treatment of both HNSCC cell lines with EGCG for 24 hours reduced the levels of ALCAM expression in a concentration-dependent manner compared to the expression in non-EGCG-treated control. These results suggest that EGCG-induced reduction in ALCAM expression is associated with an inhibitory effect of the EGCG on miR-214 expression, which may be implicated in suppressing HNSCC cell invasion.

**Figure 4:**
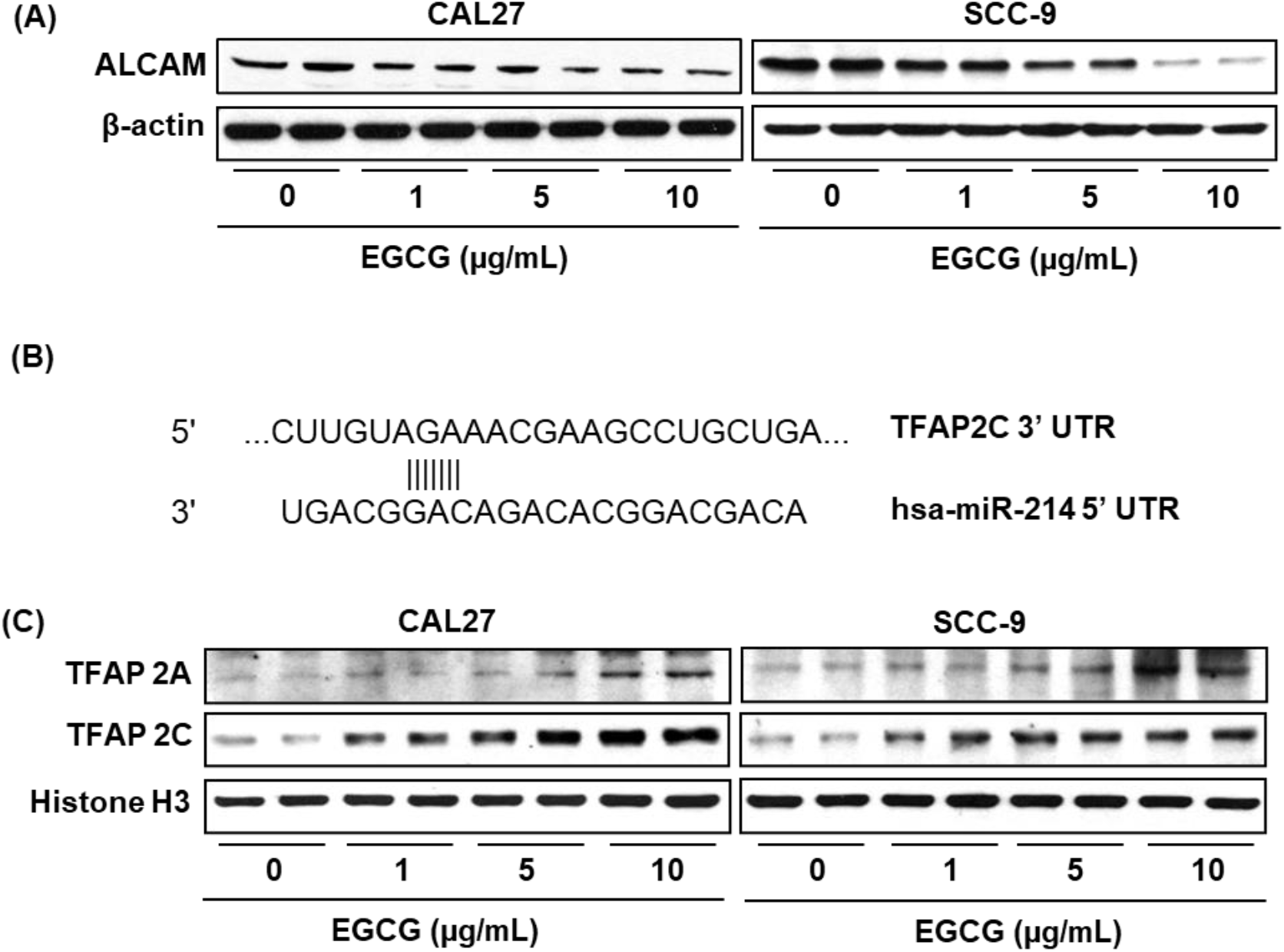
Effect of EGCG on ALCAM and TFAP2 expression in HNSCC cell lines. CAL27 and SCC-9 cells were treated with EGCG dose-dependently for 24 hours. After 24 hours of treatment, HNSCC cells were harvested, and cell lysates were prepared for western blot analysis. EGCG treatment for HNSCC cells inhibits ALCAM expression **(A)**. The binding site of miR-214 with TFAP-2C **(B)**. miR-214 regulates ALCAM expression by binding with transcription factor-activated protein 2 (TFAP2). EGCG treatment for HNSCC cells reactivates TFAP-2A and TFAP-2C proteins expression **(C)**. Each column represents an individual sample in a western blot.

### 3.5. Analysis of miR-214 binding sites with its transcription factor: In silico and western blots analysis

The transition from the non-invasive to the invasive and metastatic stage is accompanied by the loss of TFAP-2 and is associated with miR-214 overexpression. At the molecular level, to exert their functions, miRs bind with their target genes through binding in the 3’ UTR (three prime untranslated regions) and modulate their expression and function. To find the binding site of miR-214 with its regulatory gene involved in HNSCC migration/invasion, a web-based bioinformatics database TargetScan was explored to predict the target gene. According to the database, a potential binding of TFAP-2C was found in the 3’-UTR region **(Figure 4B)**. The binding of miRs with 3’ UTR of its target gene has degraded gene expression at mRNA and protein levels. The 3’ UTR, also known as silencer regions, binds to repressor proteins and inhibits the mRNA’s expression. Next, to determine whether down-regulation of miR-214 in CAL27 and SCC-9 cells following EGCG treatment alters the TFAP-2 expression, the protein levels of TFAP-2A and TFAP-2C expression in the nuclear fraction of HNSCC cell lysates were measured by western blot analysis. As shown in **Figure 4C**, EGCG treatment restores the TFAP-2A and TFAP-2C expression levels in both cell lines. These results indicated that restoration of TFAP-2 is associated with the inhibitory effect of EGCG on HNSCC invasion.

### 3.6. Dietary administration of EGCG (0.5%; W/W) inhibits the invasive capacity and establishment of tumor microenvironment in internal body organs of BALB/c nude mice

Upon metastasis, HNSCC spread throughout the body and manifested into distant organs.

The most frequent and clinical sites of HNSCC metastasis are skin, subcutaneous tissue, lungs, liver, bone, and brain[3,6]. To further verify the inhibitory effect of EGCG on HNSCC cell invading capacity, *in vivo* mouse model of metastasis was used. For this purpose, the luciferin and green fluorescent protein (GFP) tagged SCC-9 cells were injected into the tail vein of mice. Mice were imaged four weeks after the SCC-9 cells’ injection **(Figure 5A, 5B)** and sacrificed. The untreated mice (metastasis alone) showed severe metastasis as seen by multiple tumors localized in the neck region (phase contrast and luminance image). In the EGCG-fed mice, no visible tumors were seen in the neck region.

**Figure 5:**
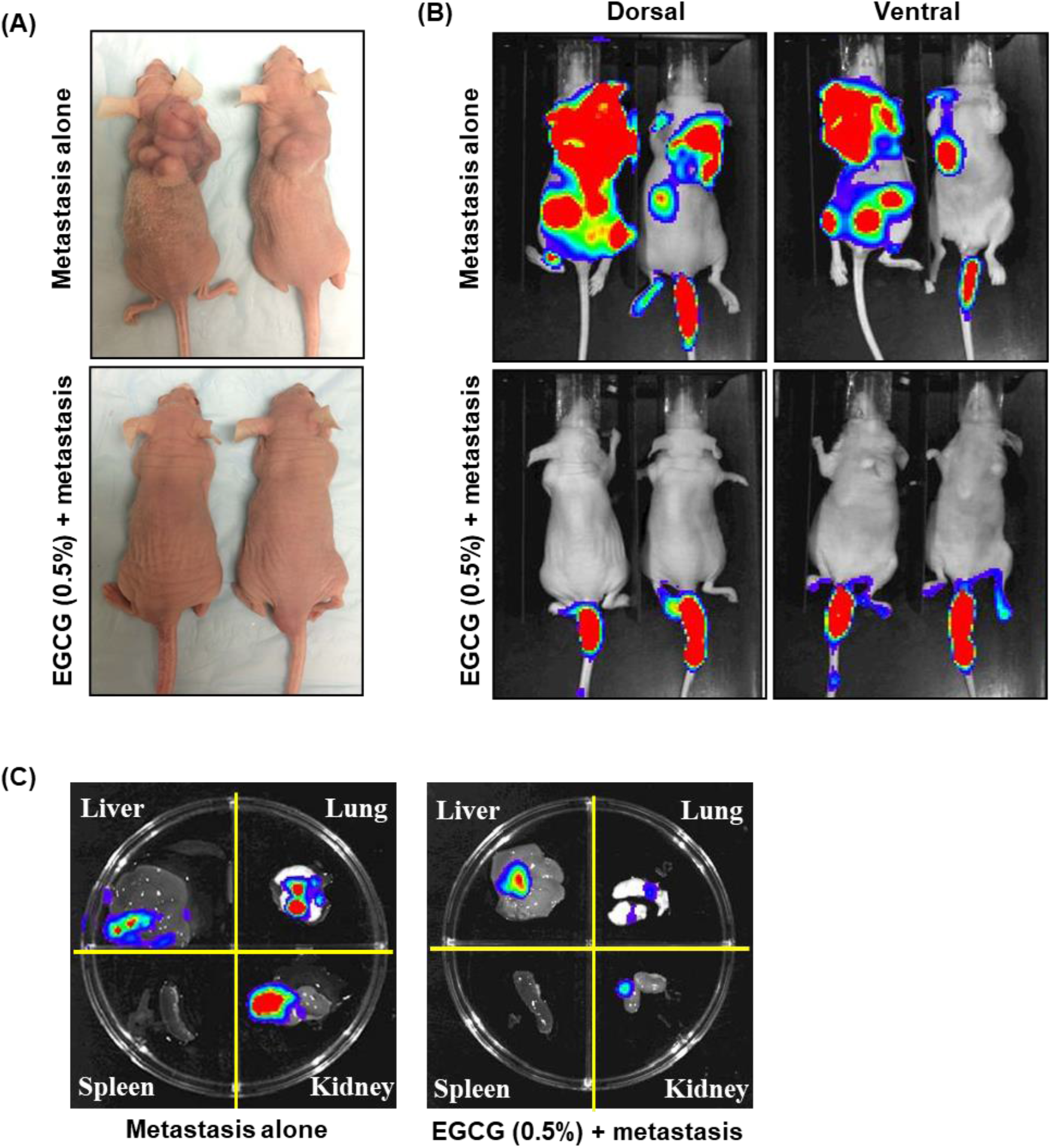
Dietary administration of EGCG inhibits the metastatic potential of SCC-9 cells in BALB/C nude mice. To facilitate the detection of experimental metastasis *in-vivo*, luciferase-tagged SCC-9 cells (2 × 10^6^) were injected into the tail vein of BALB/C nude mice with and without 0.5% EGCG treatment. After four weeks, representative bright field images of experimental mice **(A)** and luminance images of live mice **(B)** were obtained using a D-luciferin substrate. After live imaging, the mice were sacrificed, and their vital organs (liver, lungs, kidneys, and spleen) were harvested. The presence of metastatic SCC-9 cells in these organs was detected by bioluminescence imaging after spraying D-luciferin using the Xenogen IVIS200 imaging system **(C)**.

Further, to identify the distant metastasis, the mice’s vital organs, such as liver, lungs, kidneys, and spleen, were harvested and subjected to bioluminescent image analysis **(Figure 5C)**. The bioluminescence image analysis detected the presence of abundant SCC-9 cells in the liver, lungs, and kidneys, as shown by the red color (area and intensity). In contrast, a lower frequency was detected in the spleen **(Figure 5C)**. Treatment of mice with EGCG blocked invasion, accumulation, and growth of SCC-9 cells in these vital organs compared to the untreated cohort (metastasis alone), as reflected by red color intensity.

### 3.7. Dietary EGCG inhibits SCC-9 cell extravasation

For dissemination at distant sites, invasive cancer cells travel across the whole body through the circulatory system. At the favorable site of the tumor microenvironment, endothelial cells facilitate them to extravasate into another organ. As neovascularization plays an important role in cancer cell progression, endothelium-mediated invasion of cancer cells is a widely accepted cancer metastasis model. CD31, a transmembrane glycoprotein, is highly expressed in endothelium and localized at the endothelial cell junctions and plays an important role in transendothelial cellular migration. Therefore, the effect of dietary EGCG was evaluated on CD31 expression in the lungs, liver, kidneys, and spleen by immunofluorescence staining. We observed reduced CD31 expression in these organs of EGCG-fed mice compared with non-EGCG-fed (metastasis alone) mice, which ultimately leads to reduced extravasation of SCC-9 cells into these organs **(Figure 6A)**. These results suggest that EGCG administration inhibits invading capacity of HNSCC cells and blocks new vasculature formation. Tumor tissue was used as a positive control to show the presence of HNSCC cells.

**Figure 6:**
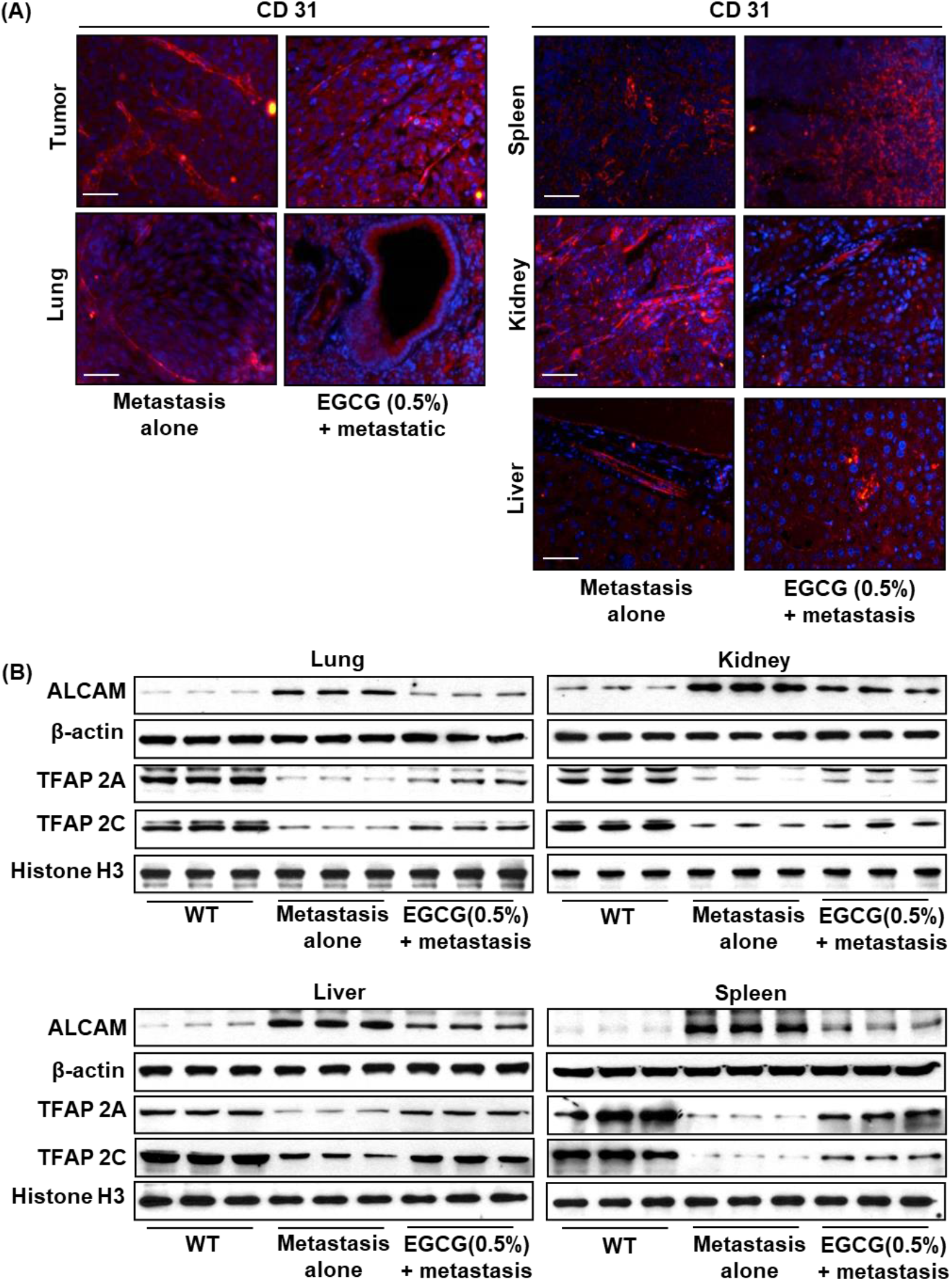
Effect of dietary EGCG on SCC-9 metastasis. The representative images of immunofluorescence staining of blood vessels by using an angiogenesis marker CD31 (red) and DAPI (blue) **(A)**. EGCG treatment in mice inhibits blood vessel formation in the vital organs of experimental mice. Effect of EGCG on ALCAM/TFAP2 proteins expression using western blot in lungs, liver, kidneys, and spleen tissues **(B)**. Each column represents an individual sample in a western blot. Scale bar = 50μm.

### 3.8. Dietary EGCG inhibits ALCAM expression and restores TFAP-2A and TFAP-2C expressions in the vital organs

Next, in continuation with our *in vivo* findings, to determine whether inhibition of invasive potential of SCC-9 cells by EGCG is associated with reduced ALCAM expression and restored TFAP-2 expression, tissues from lung, liver, kidneys, and spleen were subjected to western blot analysis. First, we compared the expression levels of ALCAM, TFAP-2A, and TFAP-2C between metastasis alone and healthy tissues (obtained from wild-type mice). The expression levels of ALCAM were higher in all four metastasis-alone cohort organs than in WT mice. In contrast, the expression of TFAP-2A and TFAP-2C was almost lost in the metastasis-alone group organs compared with the WT group. The dietary administration of EGCG (0.5%; w/w) inhibits ALCAM expression and restores TFAP-2A and TFAP-2C compared with metastatic alone **(Figure 6B)**. Our results indicated that the inhibitory effect of EGCG on the invasive potential of SCC-9 cells is mainly associated with the down-regulation of ALCAM expression and the reactivation of transcription factors (TPAF-2).

## 4. Discussion

Due to the complexity of cancer progression and the involvement of multiple factors in distant metastasis, malignant head and neck cancers are extremely difficult to treat. Although head and neck cancer primarily originates in the mucosal epithelial of the oral cavity, pharynx, and larynx, it later metastasizes in multiple organs, including the lungs, liver, kidneys, bones, brain, and skin[3,6]. The incidence of HNSCC is increasing at a very high rate and is estimated to rise upto 1.08 million per year by 2030[35]. Due to its hererogenous nature, the treatment of HNSCC remains a challenge. However, several targeted therapies have satisfactory responses at a certain point but have not achieved the desired effect. In addition to currently available interventions, treatment of HNSCC patients require aggressive multimodality approaches with the least side effect and greater efficacy in multidimensions of carcinogenesis.

The miRs are small non-coding RNAs that regulate multiple gene expressions and control cellular processes, including cell cycle, migration, invasion, and extravasation. Recent studies have reported that miRs play an important role in HNSCC disease development and progression[36-39]. The therapeutic measures that can edit miRs’ function have been recognized as an exciting new area of drug development. In addition to several nutritional benefits, bioactive molecules have been shown to have therapeutic and preventive medicinal properties and develop as a mimic or inhibitor for those miRs, which have deregulated and implicated in various diseases. Our current research aimed to reveal the potential role of miR-214 in suppressing head and neck cancer metastasis in distant organs. Deregulation of miR-214, especially over-expression, has been reported in several cancers, including but not limited to melanoma and non-melanoma skin cancer, pancreatic cancer, colorectal cancer, and nasopharyngeal carcinoma[40-44]. To investigate the potential role of miR-214 in head and neck cancer metastasis, we observed increased expression of miR-214 in four metastatic head and neck cancer cell lines used in the study, which leads to enhanced invasive characteristics. As expected, down-regulation of the miR-214 expression by either its specific inhibitor or bioactive molecule (EGCG) of green tea treatment inhibits the invasive potential of these cell lines. The invasion of metastatic cancer cells is the primary reason to make it fatal for the patients. The metastasis cascade is broadly categorized into three main processes: Invasion, intravasation, and extravasation. The invasive characteristic of cancer cells relies on the cell adhesive interactions and adhesion molecules. Studies have reported that cancer cell dissociation from its primary tumor site is accomplished by loss of function or expression of the epithelial cell adhesion molecules.

The cell adhesion molecule ALCAM has been reported to have an important role in tumor progression and is abundantly expressed in mesenchymal cells. Aberrant changes in ALCAM expression have also been reported in myeloma, prostate, esophagus, colon, pancreatic, and endometrial cancer[32,45-52]. ALCAM is also a regulator of cadherin-mediated adherens junctions in uveal melanoma cells[53]. In the ALCAM-silenced cells, the expression of mesenchymal markers (N-cadherin and β-catenin) was strikingly reduced, which dictates ALCAM’s role in the invasion of cancer cells[53]. Although the expression of ALCAM correlates with the transition of cell proliferation to invasive tumor growth and suggests its contribution to other cancers, it remains unclear in the invasive process of head and neck cancer. Earlier in melanoma, Penna et al. reported that miR-214 directly regulates ALCAM expression and plays an important role in its progression[14]. In the present study, our results demonstrated that the head and neck cancer cells also exhibit increased levels of miR-214 expression, and inhibition of miR-214 expression through EGCG treatment to head and neck cancer cells decreases ALCAM expression which eventually reduces invasion of head and neck cancer cells and metastasis observed in in-vivo and in-vitro studies. These finding suggested that EGCG mediated suppression of head and neck cancer cells’ invasion is might be due to the loss of mesenchymal property.

The changes in adhesion molecules expression are critical in carcinogenesis, and underlying molecular mechanisms responsible for these changes, which have a deleterious effect, are poorly defined. Penna et reported that the expression of ALCAM is inversely regulated by its transcriptional factors (TFAP-2)[18]. An inverse relationship between ALCAM and TFAP-2C is well documented in melanoma cells[18]. The TragetScan database indicated that miR-214 directly targets TFAP-2C through binding at 3’UTR, reduces protein level, and may regulate ALCAM expression. EGCG treatment to head and neck cancer cell lines restores TFAP-2C expression. Our in vivo findings also suggested that dietary administration of EGCG inhibits ALCAM expression. At the same time, restoring TFAP-2 expression in all vital organs inhibits invasion and extravasation of SCC-9 cells.

In conclusion, our results demonstrated that down-regulation of increased miR-214 expression in head and neck cancer cell lines through small bioactive molecule (EGCG) inhibits the invasive potential of head and neck cancer cell lines and provides new avenues to use bioactive molecules as an inhibitor of miRs to develop miR based preventive or therapeutic measures and to plan better management strategies for the distant metastasis of head and neck cancer. Further mechanistic studies need to be conducted for translational purposes.

## Abbreviations

EGCG: Epigallocatechin gallate
HNSCC: Head and neck squamous cell carcinoma
NHBE: Normal human bronchial epithelial
miR: miR
HED: Human equivalent dose
w/w: Weight by weight
TFAP2: Transcription factors AP-2 alpha
ALCAM: Activated leukocyte cell adhesion molecule

## Author Contributions

AA, VK, HF, and VKS conceived the study and participated in the design. RP provides intellectual support. AA and VK conducted all the experiments, sample collection, and analysis. All authors wrote, edited, and consented to the published version of the manuscript. The funding agency had no roles in study design, data collection, analysis, the decision to publish, or the preparation of the manuscript.

## Institutional Review Board Statement

Not applicable.

## Informed Consent Statement

Not applicable.

## Data Availability Statement

The original data presented in the study are available on request from the corresponding author.

